# Nab3 nuclear granule accumulation is driven by respiratory capacity

**DOI:** 10.1101/2022.05.23.492460

**Authors:** Katherine M. Hutchinson, Jeremy C. Hunn, Daniel Reines

**Affiliations:** Department of Biochemistry, Emory University School of Medicine, Atlanta, GA USA

**Keywords:** Nab3, granule, yeast, mitochondria, low complexity domain

## Abstract

Numerous biological processes involve proteins capable of transiently assembling into subcellular compartments necessary for cellular functions. One process is the RNA polymerase II transcription cycle which involves initiation, elongation, co-transcriptional modification of nascent RNA, and termination. The essential yeast transcription termination factor Nab3 is required for termination of small non-coding RNAs and accumulates into a compact nuclear granule upon glucose removal. Nab3 nuclear granule accumulation varies in penetrance across yeast strains and a higher Nab3 granule accumulation phenotype is associated with petite strains, suggesting a possible ATP-dependent mechanism for granule disassembly. Here, we demonstrate the uncoupling of mitochondrial oxidative phosphorylation by drug treatment or deletions of nuclear-encoded ATP synthase subunit genes, were sufficient to increase Nab3 granule accumulation and led to an inability to proliferate during prolonged glucose deprivation, which requires respiration. Additionally, by enriching for respiration competent cells from a petite-prone strain, we generated a low granule-accumulating strain from a relatively high one, providing another link between respiratory competency and Nab3 granules. Consistent with the resulting idea that ATP is involved in granule accumulation, the addition of extracellular ATP to semi-permeabilized cells was sufficient to reduce Nab3 granule accumulation. Deleting the *SKY1* gene, which encodes a kinase that phosphorylates nuclear SR repeat-containing proteins and is involved in efficient stress granule disassembly, also resulted in increased granule accumulation. This observation implicates Sky1 in Nab3 granule biogenesis. Taken together, these findings suggest there is normally an equilibrium between termination factor granule assembly and disassembly mediated by ATP-requiring nuclear machinery.

## INTRODUCTION

The RNA polymerase II transcription cycle involves RNA-binding proteins that can condense into nuclear foci to perform their functions (Boehning, et al. 2018, Cho, et al. 2018, Chong, et al. 2018, Loya, et al. 2018, Lu, et al. 2018, Sabari, et al. 2018). Proteins capable of self-assembly often contain a low complexity domain (LCD), a domain lacking stable secondary structure, that becomes structured and organized when assembling into a subcellular compartment. The LCD is often necessary for assembly and localization into a subcellular compartment.

Nab3 is an essential RNA-binding, transcription termination factor that is part of the Nrd1-Nab3-Sen1 RNA polymerase II termination complex in *Saccharomyces cerevisiae* (Arndt and Reines 2015). This termination complex is responsible for the biogenesis of snoRNAs, snRNAs, and other small non-coding RNAs such as cryptic unstable transcripts (CUTs), and serves to regulate gene expression through attenuation (Arndt and Reines 2015, Corden 2008, Dichtl 2008, Jenks, et al. 2008, Kopcewicz, et al. 2007, Kuehner and Brow 2008). Nab3 contains a well-characterized LCD that is necessary for its self-assembly into amyloid *in vitro* and its accumulation, with its dimerization partner Nrd1, into a nuclear granule *in vivo* (Alberti, et al. 2019, Carroll, et al. 2007, Darby, et al. 2012, Loya, et al. 2013, Loya, et al. 2018). The Nab3 nuclear granule shares features with cytoplasmic stress granules; both are composed of RNA-binding proteins with LCDs that can self-assemble, and both have the ability to accumulate during glucose restriction and dissolve upon refeeding (Buchan, et al. 2011, Gilks, et al. 2004, Loya, et al. 2018).

While the mechanisms responsible for the disassembly of the Nab3 granule are unknown, it has been shown that cytoplasmic stress granules can be disassembled by chaperone ATPases or DEAD-box helicases in an ATP-dependent manner (Sathyanarayanan, et al. 2020, Tauber, et al. 2020), as well as by the addition of extracellular ATP to permeabilized cells (Sathyanarayanan, et al. 2020). Taken together, sufficient respiratory capacity and proficient production of ATP are necessary for the disassembly of stress granules and by extension, may play a role in Nab3 granule disassembly as well.

The kinase Sky1 is also involved in efficient stress granule disassembly (Shattuck, et al. 2019). Sky1 is the only known conserved serine-arginine protein kinase (SRPK) expressed in *Saccharomyces cerevisiae* and has been shown to target RNA-binding proteins, aid in the disassembly of stress granules, localize to the nucleus and cytoplasm, and display a genetic interaction with Nab3, thus making it of interest as a potential protein candidate implicated in Nab3 granule disassembly (Costanzo, et al. 2016, Shattuck, et al. 2020, Shattuck, et al. 2019, Siebel, et al. 1999).

We previously developed a computational tool that facilitated the careful quantification of Nab3 granule accumulation across various commonly used laboratory strains of yeast and determined that they displayed a wide variation in the levels of granules (Hunn, et al. 2022). We considered the possibility that the variation was due in part, to differences in respiratory capacity across strains as determined by their relative abilities to grow using ethanol/glycerol as a carbon source, a commonly used method to test respiratory function in yeast. In particular, this was seen in an isolate that was a so-called *petite* strain, a commonly seen type of mutant whose mitochondrial genome is partially or completely missing. Another strain with an elevated level of granules was an S288C-based laboratory strain that is known to display mitochondrial genomic instability (Dimitrov, et al. 2009).

Here, to dissect the mechanism of Nab3 granule formation and disassembly, and explicitly test the notion that defects in respiratory capacity contribute to a high Nab3 granule accumulation phenotype, we incapacitated oxidative phosphorylation in a respiratory competent, low granule-accumulating strain through drug treatment or deletion of nuclear-encoded ATP synthase genes. Similarly, we exploited a strain with a relatively high frequency of generating petite mutants to enrich for respiratory competence and tracked changes in Nab3 granule accumulation associated therewith. We also reveal that deletion of the gene encoding the conserved serine-arginine protein kinase, *SKY1*, leads to an increase in Nab3 granule accumulation. Taken together, these data suggest a model in which Nab3 and Nrd1, presumably in concert with other factors, self-assemble into a nuclear compartment important for their function, and that ATP production supports normal levels of granule disassembly providing a mechanism to maintain an equilibrium between soluble and granule-associated transcription termination factors.

## MATERIALS AND METHODS

### Yeast Strain Construction

Yeast strains used in this paper are presented in Table 1. DY4730 was generated using high efficiency lithium acetate transformation (Gietz, et al. 1995) to transform DY4580 with a PCR product encoding *SKY1::kanMX4* amplified from plasmid pFA6a-KanMX4 (Wach, et al. 1994) using primers 5’-ACACCCCCTTTTGAGGTTGAAGAGATAGAGTAAAGAAGAAGTGTAGACATTAATGCGTACGCTGCAGGTCGAC-3’ and 5’-GAGGTTAAACAGAAAAAAAAGTAAAAGGCAAGGGCAAAATAAAGGTATAAAGGTAATCATCGACTGAATTCGAGCTCG-3’. Transformants were isolated by growth on G418 and verified by PCR.

**TABLE 1.**
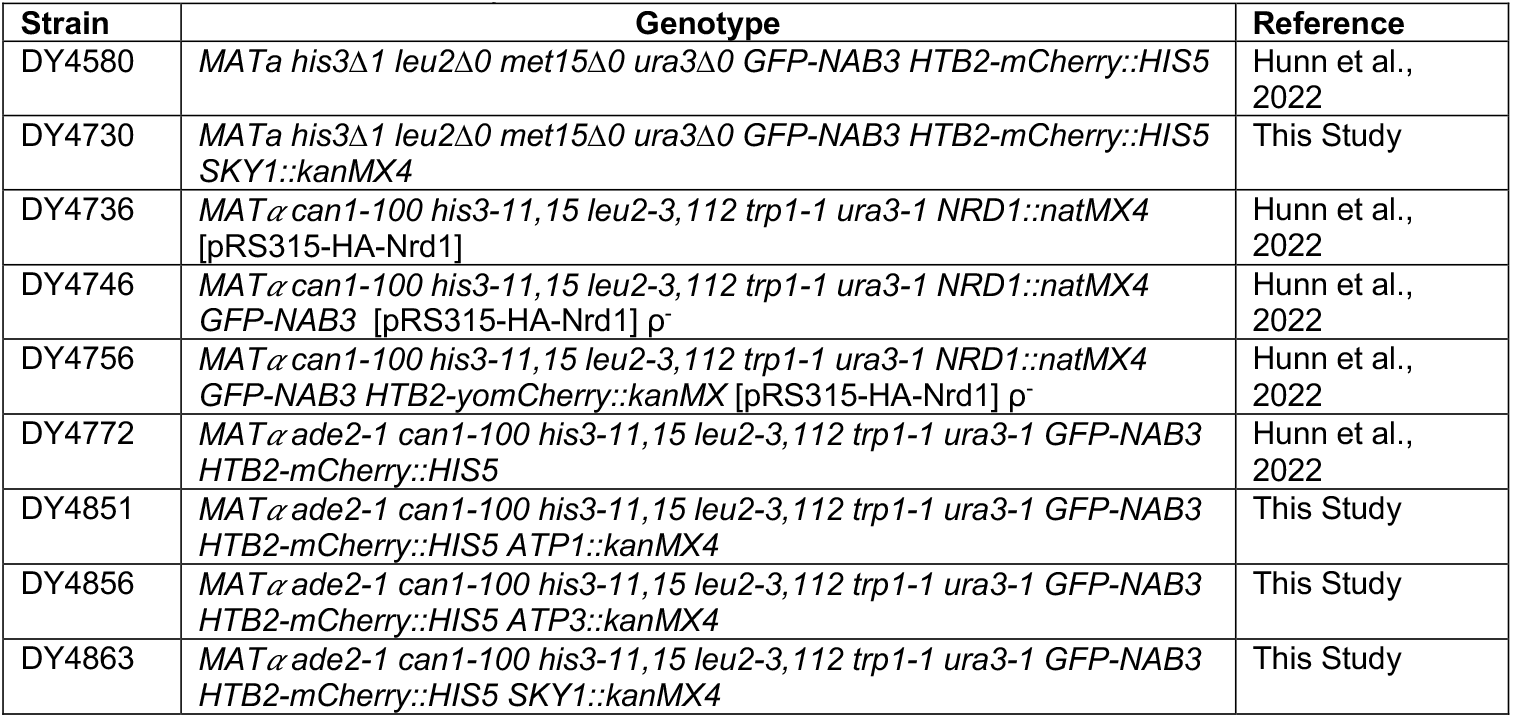
Strains used in this study.

DY4851 was generated by lithium acetate transformation of DY4772 with a PCR product encoding *ATP1::kanMX4* amplified from plasmid pFA6a-KanMX4 (Wach, et al. 1994) using primers 5’-CGCGAACCATTAGTATAACAGATTGATCGTTCAGCTCTCATACGTACGCTGCAGGT CGAC-3’ and 5’-GAGTCAGTGCTAAGAATGGAATAAACTAGAGGCTATTGTGGTCGACTGAATTCGAG CTCG. Transformants were isolated by growth on G418 and verified by PCR.

DY4856 was made as described above for DY4851. DY4772 was transformed with a PCR product encoding *ATP3::kanMX4* that was amplified using primers 5’-TACGCGAGGAACCCGGACCGCAATAACGATTAAAGAAGGCCCCGTACGCTGCAG GTCGAC-3’ and 5’-TGTTCTACAAAAACAACGTCAAATAAAGAGGCAATGCAGGGTCGACTGAATTCGAG CTCG-3’.

DY4863 was generated by lithium acetate transformation of DY4772 with a PCR product encoding *SKY1::kanMX4* that was amplified from genomic DNA of DY4730 using primer pair 5’-GTCGGGAACCGAAGCCATTG-3’ and 5’-GCAAAGTCAGCTTCCACATC-3’.

### Confocal Microscopy and Image Analysis

Confocal microscopy and image analysis was performed as described (Hunn, et al. 2022). Briefly, strains containing GFP-tagged Nab3 and fluorescently tagged histone H2B, were grown to mid-logarithmic phase, split, and washed three times into the appropriate media with or without glucose and grown at 30°C for 2 hrs. After 2 hrs, cells were pelleted and placed onto single cavity microscope slides containing 1.5% agar pads in the appropriate media. Z-stacks were captured and images were processed using FIJI (Schindelin, et al. 2012) and quantified using the MATLAB software (MATLAB. R2021a) as described (Hunn, et al. 2022). At minimum, five independent biological replicates were used for each condition and passed tests for normality. An unpaired t test with Welch’s correction was used to test for statistical significance.

For FCCP treatment, cells were grown to mid-logarithmic phase, split, and washed three times into the appropriate media with or without glucose and grown at 30°C for 2 hours. After 2 hrs, cells were treated with 20μM FCCP for 30 min. Cells were imaged as described above. For ATP treatment, cells were grown as described above for 2 hrs in the absence of glucose and treated with 0.5% DMSO for 30 minutes with or without 200 mM ATP. Cells were prepared and imaged as described above.

### Yeast Growth Assays

Yeast strains were grown to saturation at 30°C in the appropriate selection media and serially diluted for solid medium growth assays as described (Hunn, et al. 2022). Liquid growth assays were performed as described (Hunn, et al. 2022). Briefly, cultures were inoculated and grown to saturation at 30°C in the appropriate selection media. A fresh culture was inoculated using the saturated culture and grown to mid-logarithmic phase in the appropriate, glucose-containing selection media. The culture was split and washed three times into the appropriate media without glucose (starvation media) or with glucose and incubated at 30°C for 2 hours. Optical densities (600 nm) were measured, and cultures were diluted into the appropriate media to a final value of 0.05. Three 100 μL biological replicates were plated in triplicate in a 96-well plate and placed onto a BioTek Synergy H1 microplate reader at 30°C. Optical densities were recorded every 20 minutes until growth plateaued.

### Yeast Growth Assays: Selection for respiration competency

Solid medium growth assays were performed in tandem with liquid medium growth assays. First, yeast strains were grown to saturation at 30°C in the appropriate selection media. A fresh culture was inoculated using the saturated culture and grown to mid-logarithmic phase in the indicated, glucose-containing selection media. An aliquot of culture was taken, serially diluted 10-fold, and five spots of 10 μL of each dilution were applied to the appropriate solid media. Plates were grown at 30°C. An additional aliquot was taken, placed onto an agar pad, and imaged for Nab3 granule accumulation. Images were processed and analyzed as described above. The remaining culture was split, pelleted, and washed three times into the appropriate glucose-depleted media and incubated at 30°C for 2 hrs. An aliquot was taken, placed on an agar pad, and imaged for Nab3 granule accumulation. Images were processed and analyzed as described above.

For growth assays in a plate reader, optical densities were measured at 600 nm and cultures were diluted into the appropriate glucose-depleted media to a final optical density of 0.08. Three 100 μL biological replicates were plated in triplicate in a 96-well plate and placed onto a BioTek Synergy H1 microplate reader at 30°C. Optical densities were recorded every 20 min. Aliquots were taken at mid-logarithmic growth and serially diluted onto solid medium (YPD or YPEG) or placed on an agar pad and imaged for Nab3 granule accumulation. Images were processed and analyzed as described above.

### Whole Genome Sequencing Analysis

Genomic DNA for yeast strains DY4736 and DY4746 was prepared for processing at the Emory Integrated Genomics Core. A library for each strain was constructed and 150 bp paired end reads were generated on a NovaSeq 6000 S4 300 cycle flow cell to achieve 50-fold coverage. The resulting FASTQ files were uploaded to UseGalaxy.org and were mapped to the W303 RefSeq LYZE00000000 (Matheson, et al. 2017) using the sequence alignment program Minimap2 (Li 2018). The resulting mappings were visualized with the NCBI Genome Workbench, version 3.7.1 (Fig. S1).

## RESULTS

### Mitochondrial manipulations in a low granule accumulating yeast strain increase granule accumulation

Prior development of a computational tool enabled quantification of Nab3 granules which showed that accumulation of the granules in response to glucose restriction varied between yeast strains (Hunn, et al. 2022). One W303-based strain was an isolate that was extremely effective at forming Nab3 granules with ~80% [‘high’ (Hunn, et al. 2022)] of cells displaying granules. Its inability to grow when glycerol/ethanol was a carbon source suggested it was defective in respiration in contrast to its parental strain or a W303 strain (“low”) from a repository. This was further confirmed here by whole genome sequencing which revealed the high granule-forming isolate lost a substantial portion of its mitochondrial genome rendering it an authentic ρ^-^ petite, while the parental strain presented an intact genome (Fig. S1). Hunn *et al*. also observed that a low granule forming strain (~3%) treated with ethidium bromide to induce mitochondrial DNA loss, resulted in a significant increase in the fraction of cells containing granules (Hunn, et al. 2022). Since ethidium bromide treatment is a crude tool with which mitochondrial function can be manipulated, and is known to result in heterogenous effects such as an elevated frequency of nuclear DNA mutations (Rasmussen, et al. 2003), we tested the role of oxidative phosphorylation on granule accumulation with a set of more specific interventions. First, we treated a low granule accumulating strain with the mitochondrial inhibitor, carbonyl cyanide 4-(trifluoromethoxy) phenylhydrazone (FCCP) which is an ionophore that uncouples respiration from ATP synthesis by disrupting the proton gradient (Longo, et al. 1999). FCCP treatment led to a statistically significant increase in Nab3 granule accumulation in DY4772, which is an otherwise low granule accumulating strain (Figs. 1A and S2). These data strongly suggests that oxidative phosphorylation deficiencies cause an increase in Nab3 granule accumulation.

**Fig. 1.**
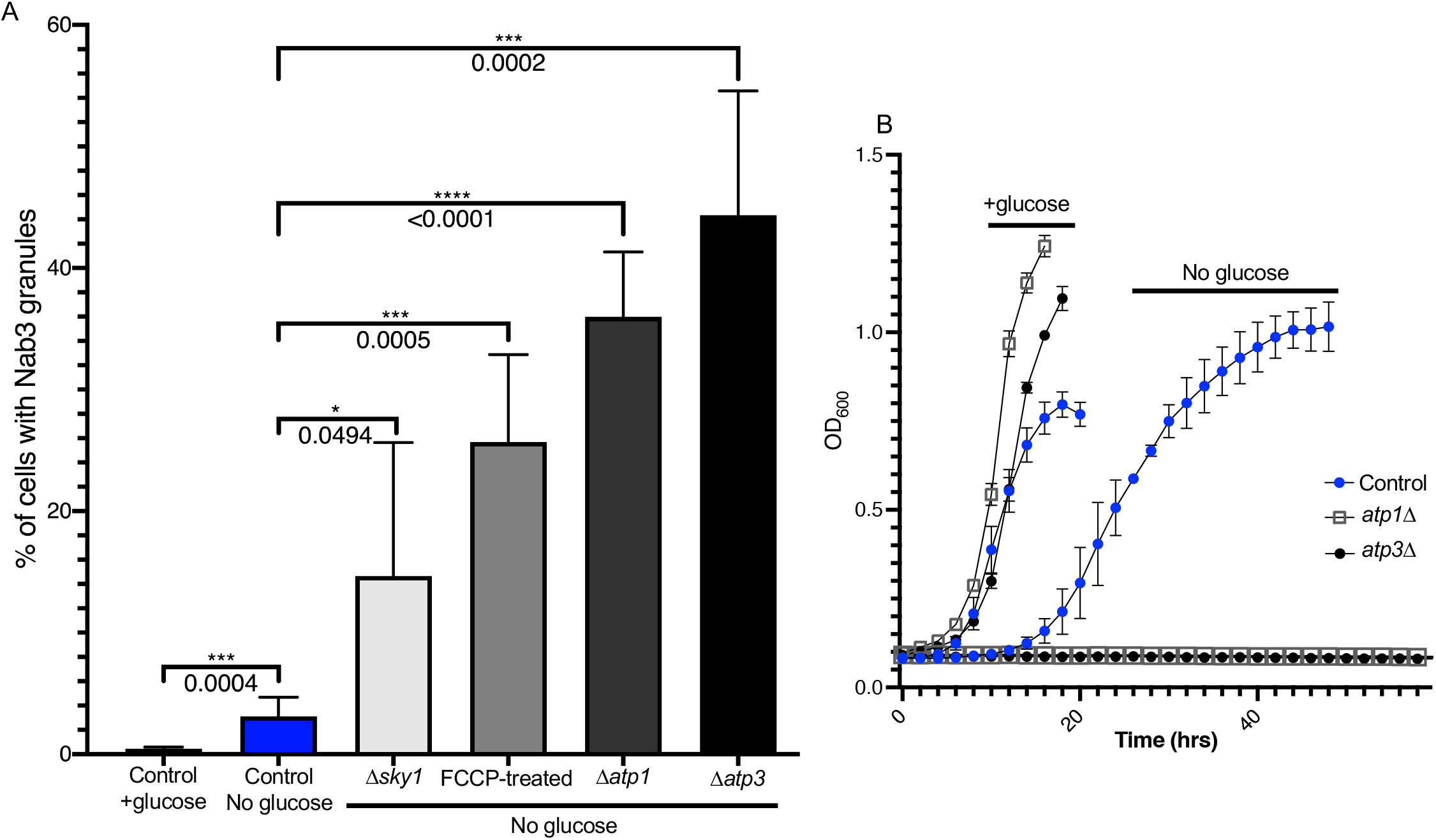
Mitochondrial manipulations in a low granule accumulating yeast strain. A. Four yeast strains (DY4772, control; DY4851, Δ*atp1;* DY4856, Δ*atp3;* DY4863, Δ*sky1*) were grown to mid-logarithmic phase, washed into starvation media, starved for 2 hours, imaged, and analyzed. Additionally, DY4772 was treated with 20μM FCCP, a mitochondrial inhibitor, for 30 minutes prior to imaging and analysis. Averages and standard deviations are plotted, and p-values are presented. [n values for the bar graph (left to right) were, 5, 10, 6, 6, 9, 6, respectively.] B. Three yeast strains [DY4772 (control, solid blue circle); DY4851 (Δ*atp1*, open square); DY4856 (Δ*atp3*, solid black circle)] were grown to mid-logarithmic phase, washed into glucose-containing media or starvation media for 2 hours at 30°C before reseeding at OD_600_= 0.08 in appropriate media. Cell growth OD_600_ was monitored at 30°C in a microplate reader. Averages and standard deviations from three technical replicates of each biological triplicate, were plotted.

Turning to a genetic approach, we independently deleted *ATP1* and *ATP3* in the low accumulating strain; these are both nuclear genes encoding subunits of the F_1_-ATPase,. Either deletion resulted in the inability of the strain to grow on ethanol/ glycerol as a carbon source (Fig. S3A) and an inability to grow upon prolonged glucose restriction (Fig. 1B), confirming that they were defective in respiration. Both strains also showed a clear increase in Nab3 granule accumulation upon glucose restriction (Fig. 1A). Taken together, these results corroborate the notion that the loss of respiratory capacity leads to an increase in the extent of Nab3 granule accumulation.

### Enrichment for a respiratory competent cell population results in a reduction of Nab3 granule accumulation

We previously discovered that the population of a commonly used laboratory strain, S288C, accumulated an intermediate amount of Nab3 granules (~50% of cells, “medium”) compared to the two strains with a low (~3%) and a high (~80%) penetrance of this phenotype (Hunn, et al. 2022). This Nab3 granule accumulating phenotype correlated with an intermediate growth phenotype on ethanol/glycerol and an intermediate ability to grow during prolonged glucose deprivation, compared to the low and high accumulating strains (Hunn, et al. 2022). The S288C lineage is known to contain mitochondrial defects and to display a high level of mitochondrial genome instability (Gaisne, et al. 1999, Shadel 1999, Young and Court 2008). These preexisting allelic differences in the S288C background contribute to a high frequency of spontaneous generation of petites with partial mitochondrial genomes (ρ^-^ cells) (Dimitrov, et al. 2009). Such cultures are mixed, containing cells with both intact and deleted mitochondrial DNA, a condition known as heteroplasmy. Since respiration is necessary for growth during prolonged glucose starvation (Weber, et al. 2020), we subjected this S288C-derived, intermediate Nab3 granule-accumulating strain to prolonged glucose starvation to enrich for a respiratory competent cell population in which petite derivatives could not survive. Cells were cultured with glucose or for an extended period (~45 hrs) of glucose deprivation, sampled at mid-logarithmic phase, and tested for their ability to grow on glycerol/ethanol (Fig. 2A). The cells from the intermediate S288C-based strain that emerged from prolonged glucose deprivation showed improved growth on glycerol (Fig. 2A, right bottom panel) as compared to the non-selected population grown in glucose medium and spotted to glycerol (Fig. 2A right top panel). Cells from both culture conditions, as well as those given the standard 2hr glucose starvation, were imaged and Nab3 granules were quantified. The relatively high level of granule accumulation observed for the 2hr starved culture was completely reversed in the prolonged starvation culture (Figs. 2B and 2C), as predicted if prolonged starvation’s selective pressure for respiratory competence reverses granule accumulation. In exploiting this “natural” variation in respiratory function found in a single culture, we obtained independent evidence again supporting the concept that Nab3 granule accumulation is driven by a loss of respiratory competence of the cell. This observation underscores the finding that heterogeneity in cell populations influences granule biology and likely other processes that are sensitive to mitochondrial function such as energy output.

**Fig. 2.**
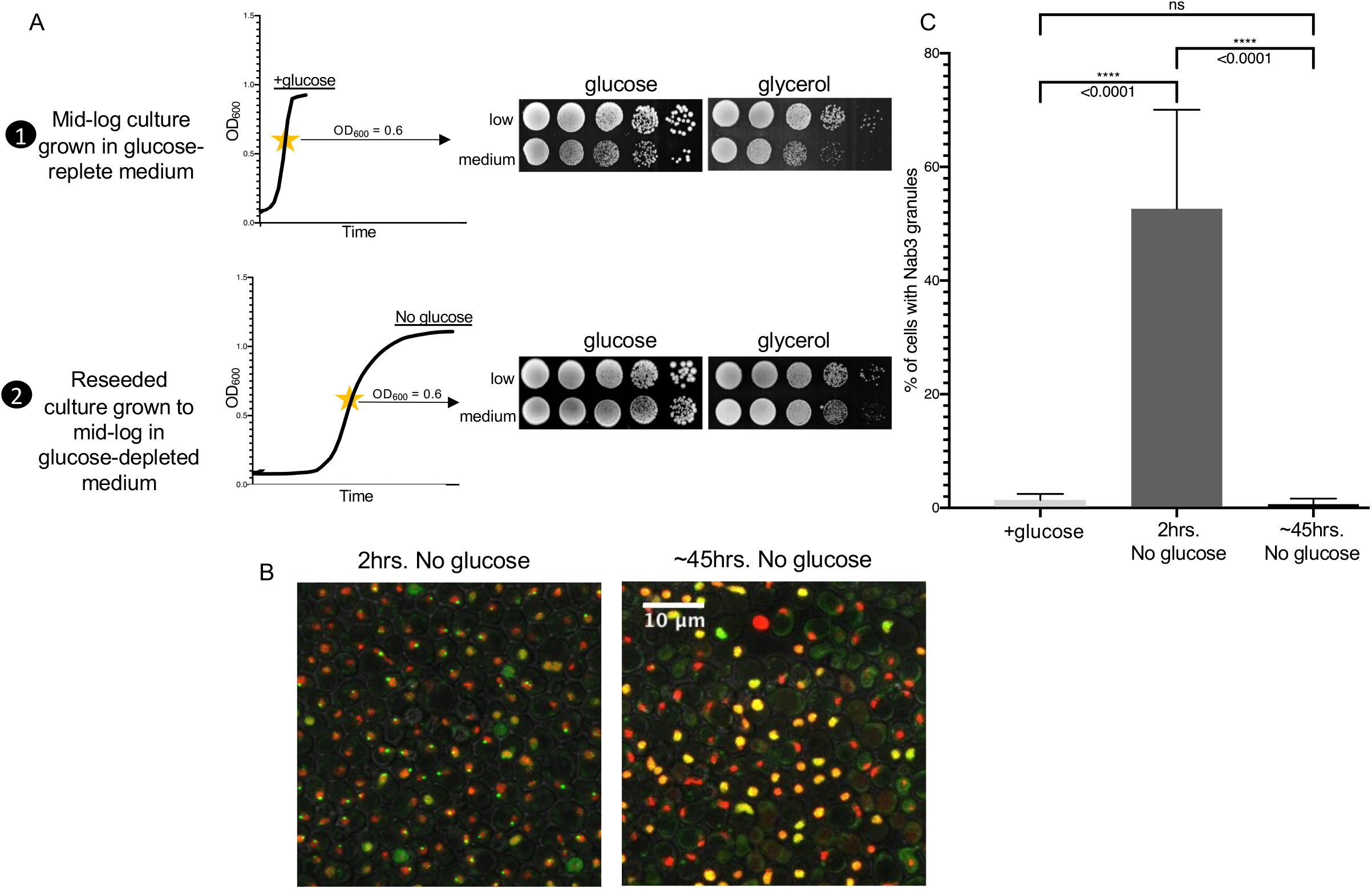
Improved respiration capacity results in a decrease in Nab3 granule accumulation. A. Yeast strains DY4772 and DY4580, which are low and medium Nab3 granule accumulating, respectively, were grown to mid-logarithmic phase in glucose, aliquots were serially diluted, and plated onto glucose (YPD) or glycerol (YPEG) solid medium (1). The remaining cells were washed into starvation media for 2 hours at 30°C before reseeding at OD_600_= 0.08. Cells were grown to mid-logarithmic phase in prolonged glucose deprivation, aliquots were serially diluted, and plated onto solid YPD or YPEG media (2). B. Yeast strain DY4580, which is medium Nab3 granule accumulating, was grown to mid-logarithmic phase, washed into starvation media, starved for 2 hrs., imaged and analyzed. At the conclusion of the 2hr. starvation, cells were reseeded at OD_600_=0.08 and grown to mid-logarithmic phase (~45hrs.) in the absence of glucose for prolonged glucose deprivation, imaged and analyzed. Representative fields of cells were imaged from independent biological replicates grown on different days. C. Medium Nab3 accumulating yeast strain DY4580 was grown to mid-logarithmic phase, washed into media with (+glucose) or without glucose, incubated for 2 hours at 30°C (2 hrs. no glucose), imaged, and analyzed. At the conclusion of the 2-hour starvation, cells were reseeded at OD_600_= 0.08 and grown to mid-logarithmic phase (~45hrs) in the absence of glucose for prolonged glucose deprivation. An aliquot was imaged and analyzed. Averages and standard deviations are plotted, and p-values are presented. [n values for the bar graph (left to right) were, 6, 29, 5, respectively. ns, not significant]

### Nab3 granule accumulation is lost by the addition of extracellular ATP

It is known that stress granule assembly, remodeling, and disassembly is modulated by ATP and that ATP is implicated in the solubilization of aggregated cellular proteins (Campos-Melo, et al. 2021, Jain, et al. 2016, Patel, et al. 2017, Takaine, et al. 2022). ATP levels drop rapidly when cells are deprived of glucose and this drop can lead to protein aggregation (Ashe, et al. 2000, Sridharan, et al. 2019, Takaine, et al. 2022, Wilson, et al. 1996). We exploited the finding that ATP added to permeabilized cells can reduce the accumulation of cytoplasmic stress granules (Sathyanarayanan, et al. 2020). Upon the addition of extracellular ATP in the presence of DMSO, the fraction of cells with a Nab3 granule were statistically significantly reduced (Figs. 3A and 3B), suggesting that ATP levels modulate Nab3 granule disassembly in a similar manner to stress granules.

**Fig. 3.**
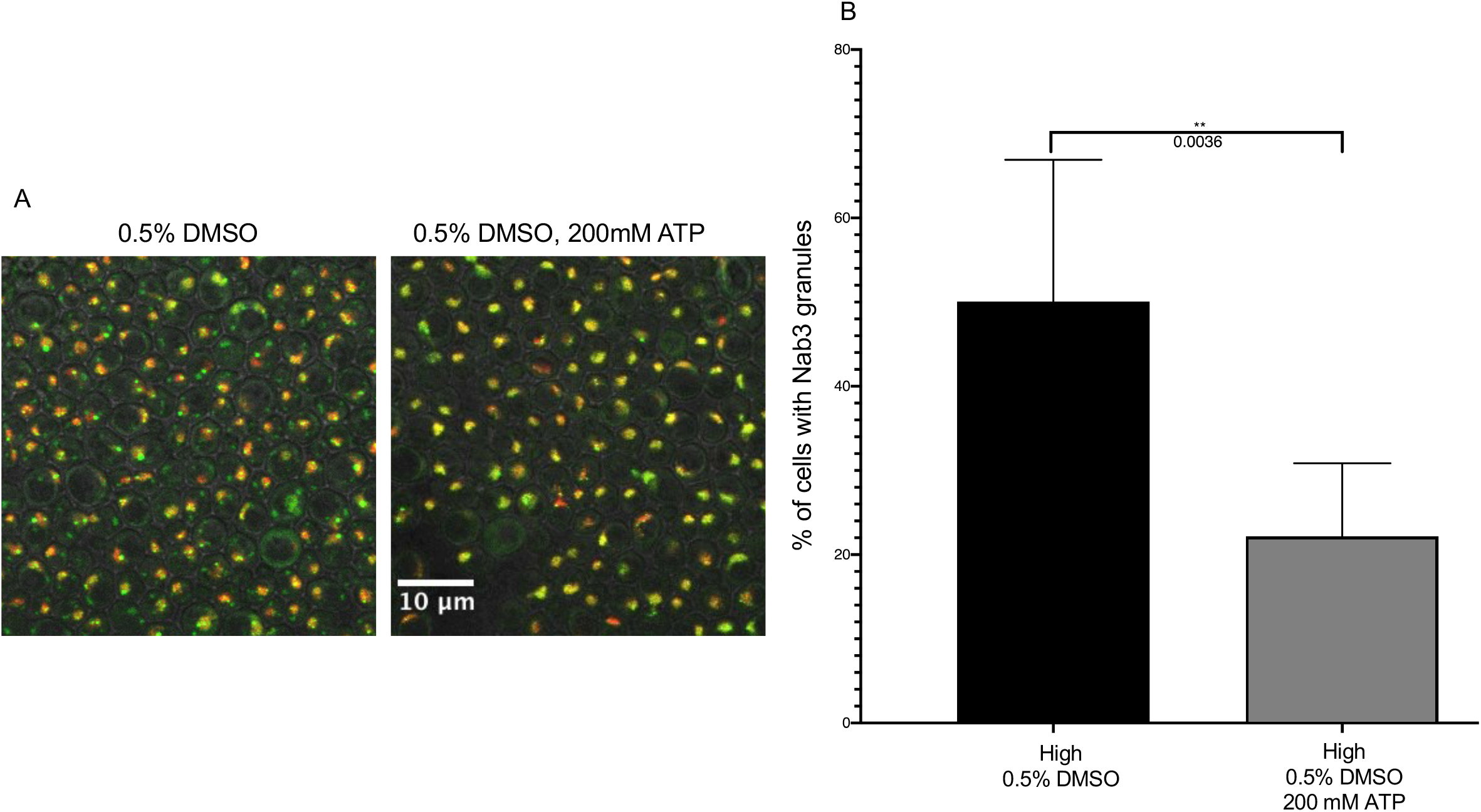
Nab3 granule levels are reduced by the addition of extracellular ATP. A. Yeast strain DY4756, which is high Nab3 granule accumulating, was grown to mid-logarithmic phase, washed into starvation media, starved for 2 hrs., treated with either 0.5% DMSO alone (control) or 0.5% DMSO and 200mM ATP, image and analyzed. Representative fields of cells were imaged from independent biological replicates grown on different days. B. Yeast strain DY4756 (‘high’) was glucose-starved for 2 hours at 30°C and treated with either 0.5% DMSO alone (control) or 0.5% DMSO and 200mM ATP for 30 min prior to analysis by fluorescent microscopy. Fields of cells were imaged from independent biological replicates grown on different days. The average and standard deviation of the percent of yeast cells containing GFP-Nab3 granules were calculated for each condition and plotted. [n values for the bar graph (left to right) were 7 and 8, respectively.]

### SKY1 knockout results in mitochondrial defects and increased Nab3 granule accumulation

Sky1 is the sole conserved serine-arginine (SR) protein kinase (SRPK) expressed in *S. cerevisiae*. It phosphorylates RNA-binding proteins, aids in the disassembly of stress granules, localizes to both the nucleus and cytoplasm, and displays a genetic interaction with Nab3 (Costanzo, et al. 2016, Shattuck, et al. 2020, Shattuck, et al. 2019, Siebel, et al. 1999). We deleted *SKY1* in the medium granule accumulating strain and found it led to a statistically significant increase in Nab3 granule accumulation (Fig. 4A). Deleting *SKY1* in our low granule accumulating strain also increased the fraction of cells with Nab3 granules (Fig. 1A). Taken together, these data implicate Sky1 in Nab3 granule biology.

**Fig. 4.**
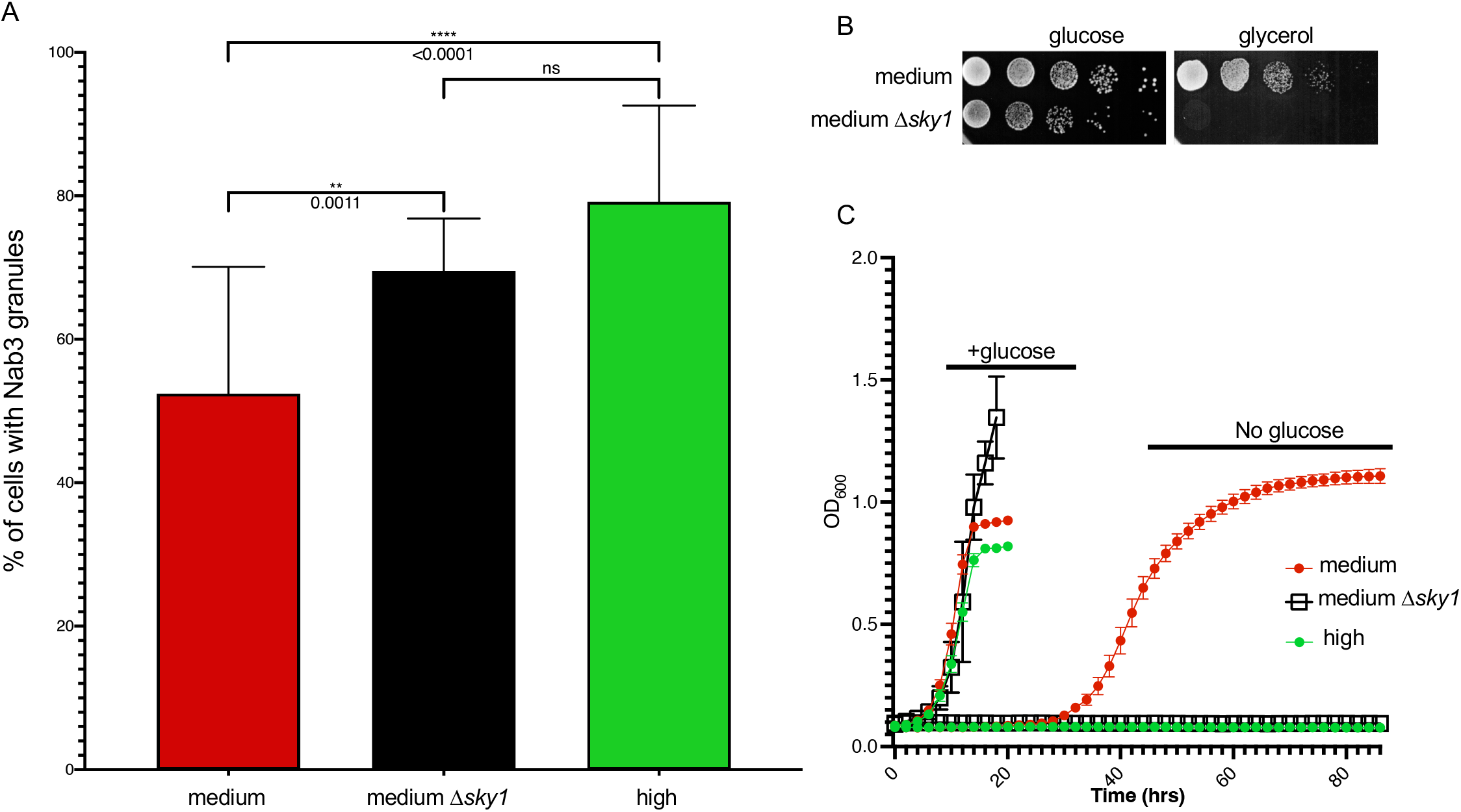
Sky1 knockout results in mitochondrial deficiency and increased Nab3 granule accumulation. A. Yeast strains DY4580 (medium), DY4730 (medium + Δ*sky1*) and DY4756 (high) were glucose-starved for 2 hrs at 30°C and analyzed by fluorescent microscopy. Fields of cells were imaged from independent biological replicates grown on different days. The average and standard deviation of the percent of yeast cells containing GFP-Nab3 granules were calculated for each condition and plotted. [n values for the bar graph (left to right) were 28, 6, 10, respectively.] B. Yeast strains DY4580 (medium) and DY4730 (medium + Δ*sky1*) were grown to saturation and diluted with sterile water to a concentration of 10^7^ cells/mL. Cells were serially diluted 10-fold and 10 μL spots of each dilution were spotted on SC-glucose or SC-glycerol/ethanol as indicated and incubated at 30°C. C. Three different yeast strains (DY4580 (medium, solid red circle); DY4730 (Δ*sky1*, open black square), DY4756 (high, solid green circle) were grown to mid-logarithmic phase, washed into glucose-containing media or starvation media for 2 hrs at 30°C and reseeded at OD_600_= 0.08 in appropriate media. Growth of cells was monitored at 30°C in a microplate reader by measuring OD_600_ (600 nm). Averages and standard deviations from three technical replicates of each biological triplicate, were plotted.

Since respiration defects lead to an increase in granule accumulation (Figs. 1-3), we tested if these strains also had a respiration defect. Both Sky1 deletion strains displayed impaired respiration compared to their parental strains (Figs. 4B, S3) with the S288C-based strain suffering a severe, and perhaps complete, inability to grow on glycerol/ethanol. Regardless of strain background, it is clear that the loss of *SKY1* resulted in a mitochondrial defect. In a further test of respiratory competence, prolonged growth in the absence of glucose was completely impaired when the S288C strain’s *SKY1* gene was deleted, effectively converting the growth pattern of a lower granule-accumulating strain into the growth pattern of a petite strain that accumulates high levels of granules (Fig. 4C). Thus, the Sky1 role in Nab3 granule biology appears to be mediated by its mitochondrial function.

## DISCUSSION

Here we provide multiple lines of evidence that strongly implicate mitochondria and respiration in the biogenesis of Nab3 granules. This establishes the first documentation for the underlying regulatory mechanism of Nab3 granules, which resemble stress granules. Nab3 and Nrd1, are, to our knowledge, the only transcription termination factors for which conditional assembly into a granule has been reported. Whether the granule is a complex ribonucleoprotein structure that is an active termination factor, or simply an aggregate formed from the Nab3 and Nrd1 proteins that both harbor prion-like domains, remains to be elucidated.

The relationship between Nab3 granules and respiration can be understood through a model (Fig. 5) in which there is normally an equilibrium between pan-nuclear (soluble) Nab3 and granule-associated Nab3 due to the action of an ATP-requiring granule dissociation machinery. In the presence of glucose (Fig. 5, condition 1), Nab3 generally localizes throughout the nucleus when ATP generation using glycolysis is unencumbered and Nab3 condensation is checked by disassembly-ATPases. If respiration is compromised (Fig. 5, condition 2), the removal of glucose leads to reduced ATP levels, the disassembly machinery cannot keep Nab3 soluble, and it accumulates into a nuclear granule. In periods of prolonged glucose deprivation (Fig. 5, condition 3), cells capable of respiration generate ATP and proliferate, resulting in the disassembly of the Nab3 granule and a return to a pan-nuclear distribution. Note that a small but statistically significant subset of cells display granules even when cells are fully respiration competent (Fig. 1, leftmost two bars), contributing to the idea that there is a constant equilibrium that shifts toward assembly when ATP levels become limiting (Hunn, et al. 2022).

**Fig. 5.**
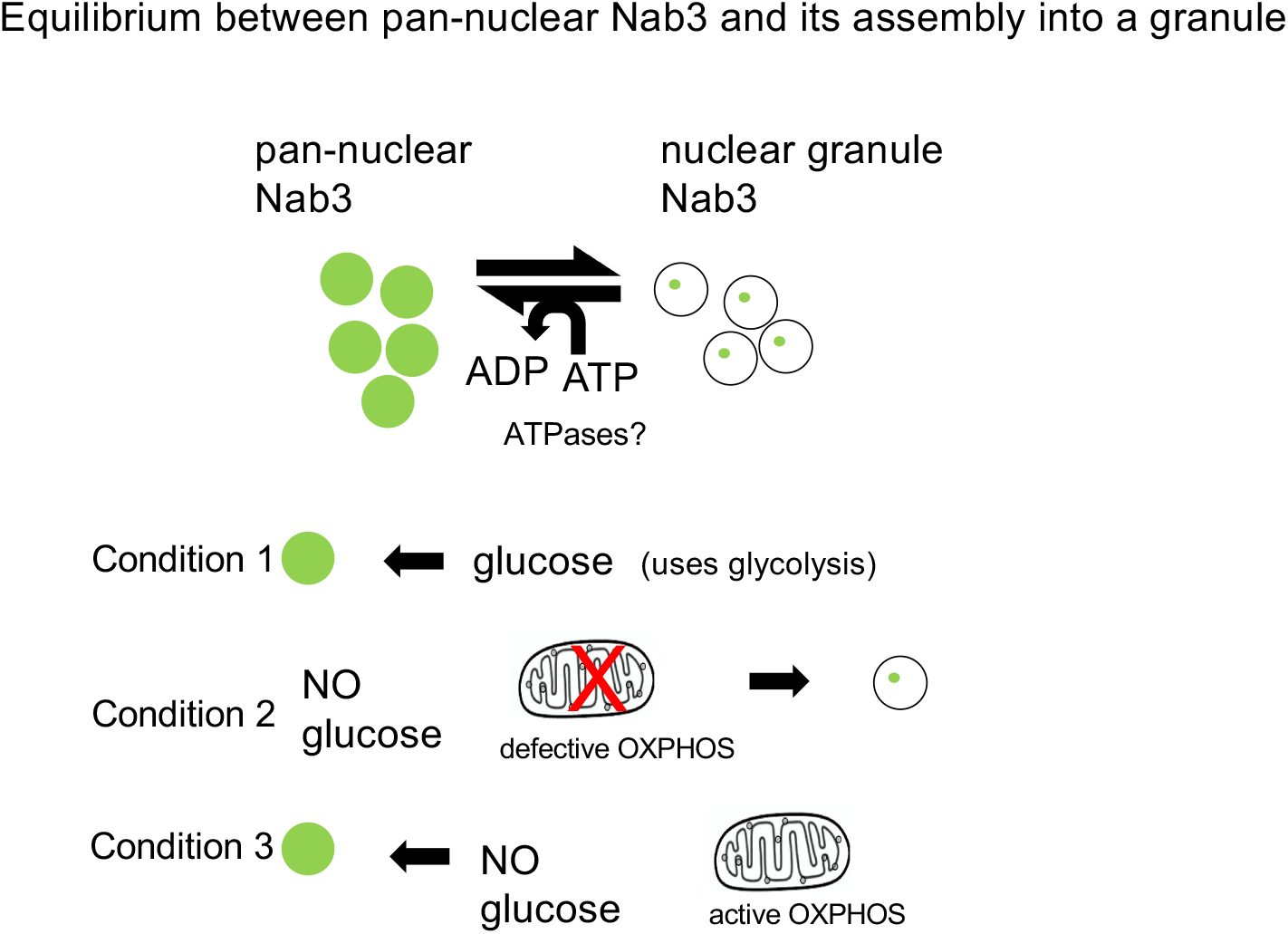
Equilibrium between pan-nuclear Nab3 and its assembly into a nuclear granule: a working model for Nab3 granule disassembly and accumulation. There is an equilibrium between pan-nuclear and granule-associated Nab3. In the presence of glucose (condition 1), Nab3 generally localizes throughout the nucleus when ATP generation using glycolysis is unencumbered and Nab3 condensation is reversed. If respiration is compromised (“X”, condition 2), the removal of glucose reduces ATP levels, the disassembly machinery cannot keep Nab3 soluble, and the protein accumulates into a nuclear granule. In periods of prolonged glucose deprivation (condition 3), cells capable of respiration can generate ATP and proliferate, resulting in the disassembly of the Nab3 granule and a return to a pan-nuclear distribution.

Precedent for this model can be found from yeast stress granules, which are dispersed by the ATP-dependent dissociation activity of the Hsp104 chaperone ATPase (Sathyanarayanan, et al. 2020). Other ATP-dependent mechanisms that regulate protein condensation and stress granule abundance have been identified including DEAD box ATPases (Weis and Hondele 2022) and AAA ATPases (Buchan, et al. 2013). Alternatively, high concentrations of ATP may act as a hydrotrope in which it is not hydrolyzed but acts as an amphiphile to directly solubilize phase separated protein condensates (Patel, et al. 2017). In either case, it follows that treatments, challenges, or genetic variations that compromise ATP production limit the dissociation ability of this machinery and lead to granule accumulation because they cannot be efficiently dissolved. The extent to which cells in the population accumulate granules would be a function of the severity of the loss in energy generation. As would be expected, abrupt reversibility of Nab3 granules becomes obvious when cells are refed glucose (Loya, et al. 2018).

Variation in the extent to which different strains spawn daughter cells that have a partial or complete absence of the mitochondrial genome (ρ^-^ or p^0^ petites), and as a result, manifest a reduced respiratory capacity, yields strain to strain differences in granule count. For some genetic polymorphisms, such as that seen for mitochondrial DNA polymerase gamma *MIP1*, it is easy to grasp how a sequence variant can increase the frequency of mitochondrial genome loss (Young and Court 2008). Indeed, variants in the sequence of the conserved *MIP1* human ortholog are linked to mitochondrial disease (Baruffini, et al. 2006, Kaliszewska, et al. 2015). For others, such as the experimentally generated *Dsky1* mutation reported here, the mechanistic relationship between the gene product’s function and loss of respiratory capacity needs to be studied further.

A high throughput screen of a collection of yeast gene deletion strains showed the *SKY1* deletion has reduced ability to grow on ethanol as a carbon source, implicating Sky1 in mitochondrial function (Tigano, et al. 2015). Our findings confirm and extend the relationship between Sky1 and mitochondria and suggests its role in influencing the assembly state of both cytoplasmic stress granules (Shattuck, et al. 2019) and nuclear Nab3 granules, could be in providing ATP for the dissociation machinery of each compartment. This explanation is supported by a genetic interaction between *SKY1* and *HSP104* (Shattuck, et al. 2019) Interestingly, a global genetic interaction survey also picked up a *NAB3-SKY1* synthetic genetic effect which deserves more investigation (Costanzo, et al. 2016). Alternatively, the Sky1 kinase could act more directly in each compartment, perhaps by phosphorylating a granule component, as it is known to shuttle between the nucleus and cytoplasm (Shattuck, et al. 2020). Mutation of enzymes involved in biosynthesis of the adenine ring or phosphorylation of the nucleotide to the triphosphate level, also modify the ATP energy charge of cells in a manner that can lead to protein aggregation in yeast (Takaine, et al. 2022).

This report extends to a nuclear transcription termination complex known to undergo regulated compaction; the principles of compartment assembly observed for stress granules. These findings should provide clues to the elucidation of the mechanism of assembly and disassembly of this essential complex and the consequence the resulting compartmentalization of the complex has upon RNA metabolism.

---

**Fig. S1.**
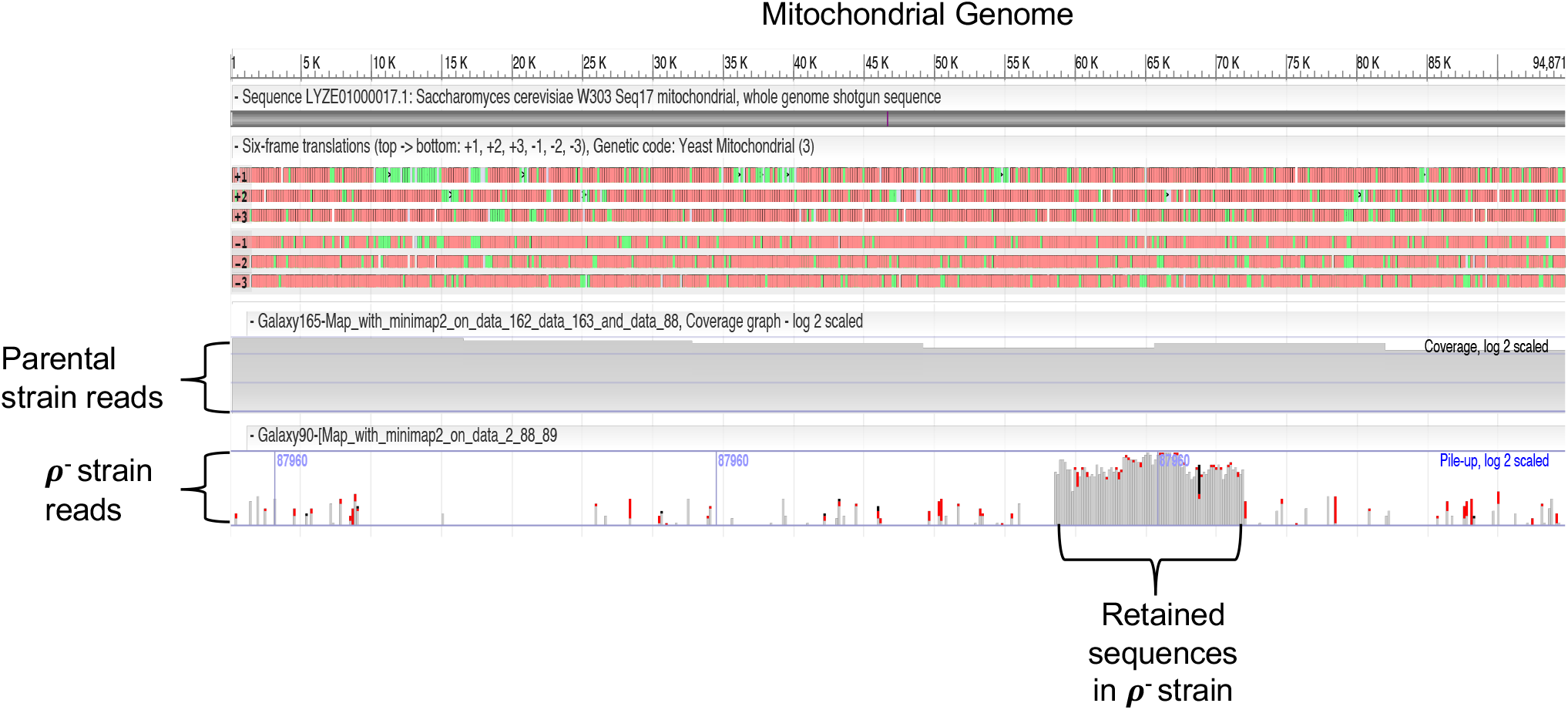
A high granule accumulating yeast strain is ρ^-^. Whole genome sequencing analysis reveals that the high granule accumulating yeast strain DY4746 (parent to DY4756) is ρ^-^ as it possesses only a small portion of its mitochondrial genome compared to its parental strain (DY4736). Sequencing reads in the lower two panels of the NCBI viewer are piled in horizontal lines with gray representing identities to the reference sequence and mismatches shown in red. All but ~15 kbp of mitochondrial DNA are lost from this petite strain.

**Fig. S2.**
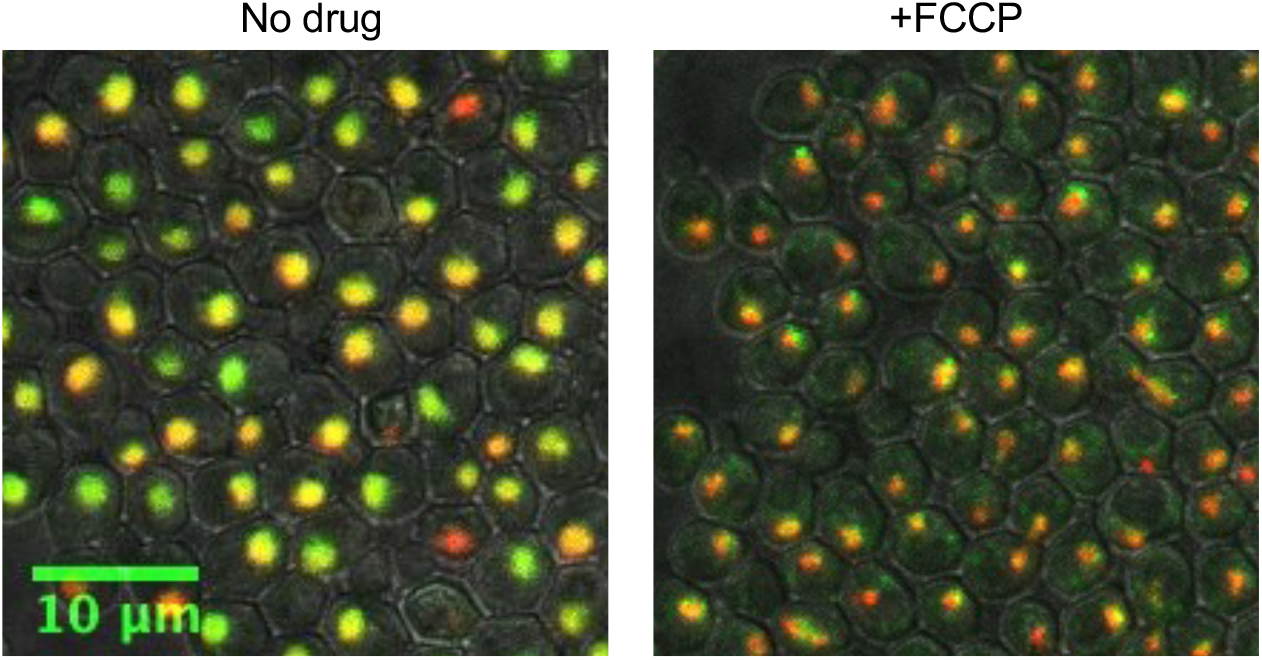
Confocal microscopy of a glucose starved low granule accumulating strain treated with and without the electron transport uncoupler, FCCP. Yeast strain DY4772, which is low Nab3 granule accumulating, was grown to mid-logarithmic phase, washed into starvation media, starved for 2 hrs, treated with mock or 20μM FCCP for 30 min, imaged, and analyzed.

**Fig. S3.**
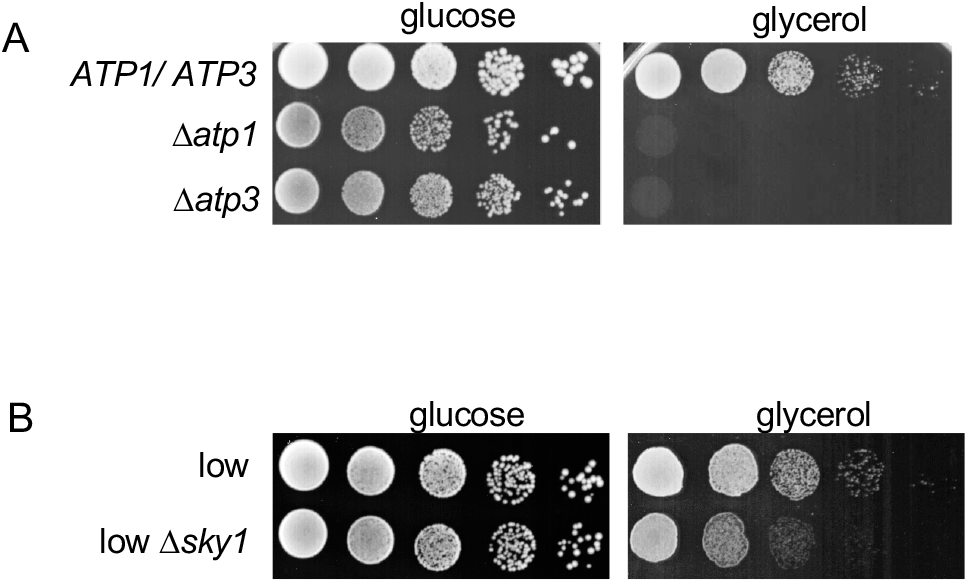
Deletions of nuclear encoded genes result in a mitochondrial defect. A. Yeast strains DY4772 (control), DY4851 (*Datp1*) and DY4856 (*Datp3*) were grown to saturation and diluted with sterile water to a concentration of 10^7^ cells/mL. Cells were serially diluted 10-fold and 10μl of each dilution were spotted onto SC-glucose or SC-glycerol/ethanol as indicated and incubated at 30°C. B. Yeast strains DY4772 (control) and DY4863 (low Δ*sky1*) were grown to saturation and diluted with sterile water to a concentration of 10^7^ cells/mL. Cells were serially diluted 10-fold and spotted on SC-glucose or SC-glycerol/ethanol as indicated and incubated at 30°C.

## Statements and Declarations

### Funding

This work was funded by National Institutes of Health (R01 GM120271 to D.R.), the Emory University School of Medicine, the Emory University Research Committee, and the Emory Integrated Cellular Imaging Core. The content is solely the responsibility of the authors and does not necessarily reflect the official views of the National Institute of Health.

### Competing Interests

The authors have no relevant financial or non-financial interests to disclose.

### Author Contributions

All authors contributed to the study conception and design. Material preparation, data collection and analysis were performed by Katherine Hutchinson, Jeremy Hunn, and Daniel Reines. The first draft of the manuscript was written by all authors and previous versions of the manuscript were commented upon by all authors. All authors read and approved the final manuscript.

### Data Availability

All data generated or analyzed during this study are included in this published article [and its supplementary information files].

### Ethics Approval, Consent to Participate, Consent to Publish

No human subjects or vertebrate animals were used in this work.

## Notes

### Competing Interest Statement

The authors have declared no competing interest.

